# Disruption of innate defense responses by endoglycosidase HPSE promotes cell survival

**DOI:** 10.1101/2020.08.05.238758

**Authors:** Alex Agelidis, Benjamin A. Turturice, Rahul K. Suryawanshi, Tejabhiram Yadavalli, Dinesh Jaishankar, Joshua Ames, James Hopkins, Lulia Koujah, Chandrashekhar D. Patil, Satvik R. Hadigal, Evan J. Kyzar, Anaamika Campeau, Jacob M. Wozniak, David J. Gonzalez, Israel Vlodavsky, Jin-ping Li, David L. Perkins, Patricia W. Finn, Deepak Shukla

## Abstract

The drive to withstand environmental stresses and defend against invasion is a universal trait extant in all forms of life. While numerous canonical signaling cascades have been characterized in detail, it remains unclear how these pathways interface to generate coordinated responses to diverse stimuli. To dissect these connections, we follow heparanase (HPSE), a protein best known for its endoglycosidic activity at the extracellular matrix but recently recognized to drive various forms of late stage disease through unknown mechanisms. Using herpes simplex virus-1 (HSV-1) infection as a model cellular perturbation, we demonstrate that HPSE acts beyond its established enzymatic role to restrict multiple forms of cell-intrinsic defense and facilitate host cell reprogramming by the invading pathogen. We reveal that cells devoid of HPSE are innately resistant to infection and counteract viral takeover through multiple amplified defense mechanisms. With a unique grasp of the fundamental processes of transcriptional regulation and cell death, HPSE represents a potent cellular intersection with broad therapeutic potential.

## Introduction

As major constituents of the extracellular matrix of virtually all cells, heparan sulfate and other protein-associated glycans have been well studied as co-receptors for a variety of macromolecules and pathogens. However, very little understanding of their regulatory mechanisms in cell signaling, microbial invasion, and cellular physiology exists. Some recent studies have made startling observations that enzymatic splitting of these sugars at the plasma as well as nuclear membranes may constitute an important regulatory step that promotes inflammation and tissue invasion (1-3). An endoglycosidase, heparanase (HPSE), potentially contributes to pathological inflammatory signaling through its glycosidic action on the extracellular matrix. HPSE is the only known mammalian enzyme capable of splitting polymers of heparan sulfate, and heparan sulfate is the only known enzymatic target of HPSE (1). These chains of heparan sulfate hydrolyzed by HPSE may be present in multiple cellular locations in the context of various proteoglycans (4). Yet HPSE remains a mysterious cellular component, as additional reports of its tendency to appear in the nucleus and regulate gene expression continue to emerge (5, 6). Associations between HPSE and various pathologies, including inflammation and cancer metastasis, have historically been attributed to the protein’s cleavage of HS chains at the cell surface and basement membrane (1, 2); however, HPSE may possess unique roles arising independently of its enzymatic active site. Recently our group and others have described yet another important role for HPSE as a driver of microbial pathogenesis. HPSE is now known to be transcriptionally upregulated by several viruses dependent upon heparan sulfate for attachment and entry, and subsequently facilitates egress of newly produced viral particles (7-10). While enzymatic activity of HPSE is believed to enable viral release through splitting of HS residues at the cell surface, it has been reported to also regulate gene expression and trigger proviral signaling through some distinct non-enzymatic activity (11). With a focus on cellular responses to infection, we show here that HPSE acts beyond its known endoglycosidase activity as a potent regulator of the signal transduction phase of cellular defense.

## Results

### Cells lacking HPSE are intrinsically resistant to HSV-1 infection

Given our unexpected finding that HPSE can directly drive viral pathogenesis (7, 11), we became interested in investigating cellular responses to infection in the absence of HPSE altogether. To our further surprise, HSV-1 infection of Hpse-knockout (KO) mouse embryonic fibroblasts (MEFs) showed a dramatic reduction in virus production, as compared to wildtype MEFs. Virus enters cells at normal levels, but viral replication and protein production are decreased by several orders of magnitude **(Figures 1 and S1)**. Hpse-KO cells originally appear capable of immediate early viral gene production, but late gene products are effectively absent **(Figure 1A)**. These findings align with previous work from our group and others’ showing that active HPSE translocates to the nucleus and influences gene expression through an unknown mechanism (5, 6, 12-14). We therefore adopted an unbiased approach to generate a clearer understanding of how HPSE, or lack thereof, potently influences gene expression and cell signaling to restrict viral production. RNA-seq analysis constituted the primary exploratory and gene clustering pipeline, while quantitative proteomics analysis of the same samples provided an additional level of functional validation. The transcriptomic landscape of Hpse-deficient cells is markedly altered by ablation of this single locus, with 1,385 genes showing baseline differences in expression **(Figure S1)**. Based on gene ontology (GO) analysis of non-infected cells, we observed that Hpse-deficient cells show significant enrichments of genes representing pathways of “defense response to virus” and “activation of immune response”, suggesting that these cells are somehow intrinsically resistant to infection **(Figure 1D)**. As an independent confirmation, differential expression analysis of the only existing publicly available dataset of HPSE genetic alteration (GSE34080) yielded strikingly similar results **(Figure S2)**. This silencing RNA knockdown of HPSE performed by other investigators in human gastric cancer cells shows significant enrichments of GO terms including “negative regulation of viral genome replication” and “response to type I interferon”, indicating a broad external validity of these newly described HPSE functions.

**Figure 1.**
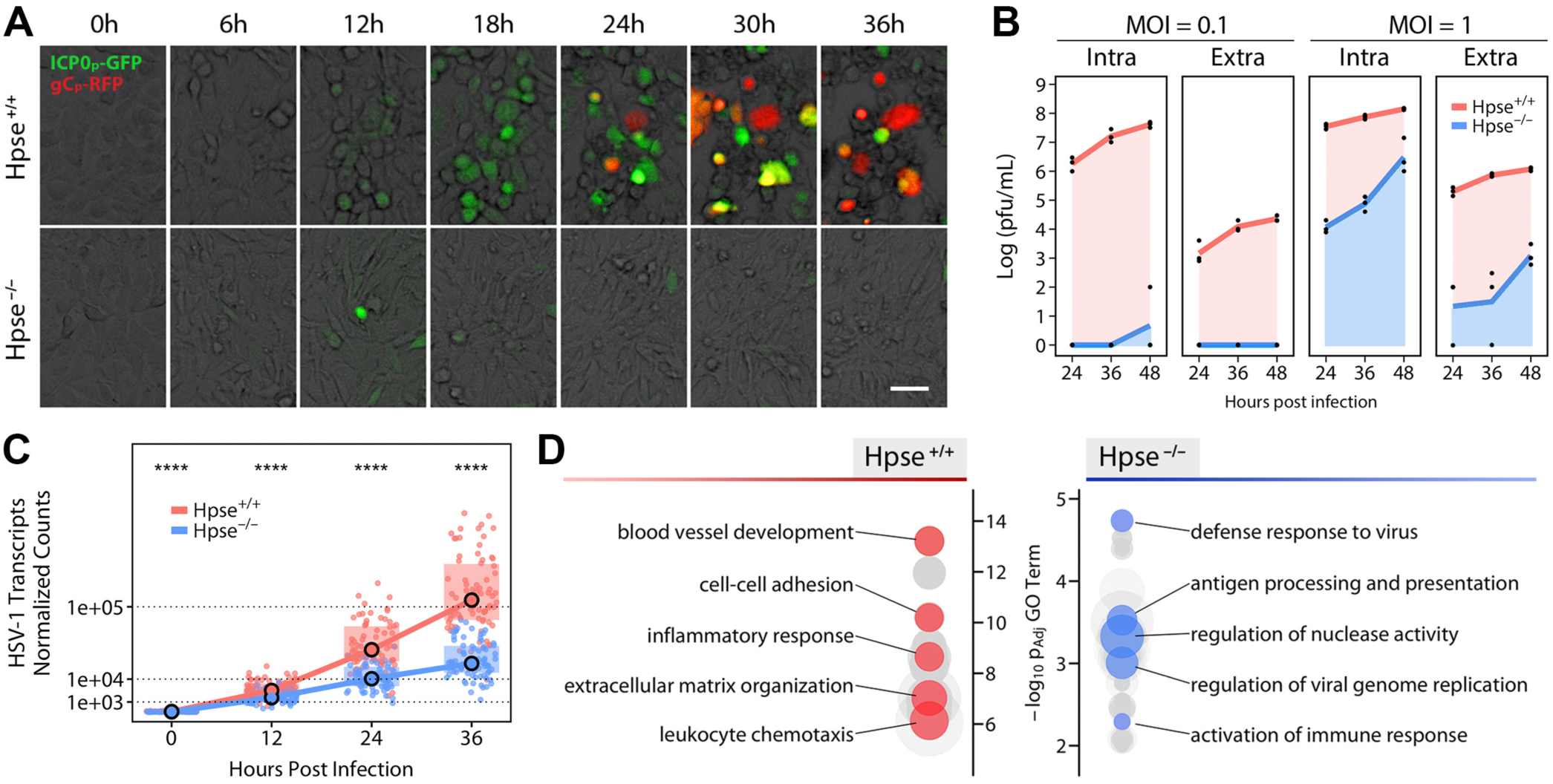
Baseline differences in host gene expression grant HPSE-deficient cells resistance to viral infection. **(A)** Representative images of infection timecourse of wildtype (Hpse ^+/+^) and heparanase-deficient (Hpse ^-/-^) mouse embryonic fibroblasts (MEFs) with a dual fluorescence strain of HSV-1. Green signal indicates viral immediate early gene expression (ICP0) while red signal indicates viral late gene expression (gC). Scale bar, 50 μm. **(B)** Titers of intracellular (Intra) and extracellular (Extra) HSV-1 detected at two multiplicities of infection (MOI) in wildtype and Hpse ^-/-^ MEFs (*n*=3). **(C)** HSV-1 viral gene expression as quantified by transcriptomics analysis. Each data point represents one of 74 detected viral transcripts. Median and interquartile range are plotted, significance analyzed by Wilcoxon signed-rank test. **(D)** Baseline differential enrichment of gene ontology terms of Hpse ^+/+^ and Hpse ^-/-^ cells in the absence of infection. Each node represents a unique term, with node size representing fold enrichment. Select summary terms are highlighted. ****p<0.0001. See also Figures S1 and S2.

### Temporal viromics catalogs transcriptional landscape shifts dependent on HPSE

Hundreds of gene expression changes occur in a viral infection, and as such, viruses represent excellent tools to probe the connections between cellular processes and signaling cascades. To appreciate major regulators of this cellular remodeling and understand how these shifts are influenced by HPSE, we generated clusters of the significantly differentially expressed genes based on temporal expression patterns. Topological clustering using a t-distributed stochastic neighbor embedding (t-SNE) (15) followed by an affinity propagation (APcluster) (16) algorithm produced 13 well-defined clusters **(Figures 2A, S3 and S4)**. Further attention to the most heavily induced clusters 9 and 12 revealed that Hpse-deficient cells are hyperactive in antiviral cytokine signaling and cellular defense, while infection of wildtype cells stimulates production of cellular machinery essential for virus assembly **(Figures 2B and 2C)**. In fact, cluster 12 also contains all viral genes, which are transcribed virtually unimpeded in wildtype cells, while expression of this cluster is stalled in Hpse-KO, concordant with the defense response induction occurring around 12 hpi. We then scanned upstream promoter regions of genes in each cluster using the PASTAA algorithm to identify major transcriptional regulators and correlate with temporal expression patterns **(Figure 2D)**. Our finding of highly significant enrichments of multiple interferon regulatory factors (IRFs) and nuclear factor (NF)-κB as the primary regulators of cluster 9 expression adds further credence to results of the above GO analysis and previous *in vitro* evidence of an inverse relationship between IRF7 and HPSE gene expression (11). Interestingly, this analysis also designated cAMP responsive element binding protein (CREB) as a major proviral factor, and remarkably, pharmacological blockade of various components of the CREB signaling interface resulted in potent inhibition of virus production in both wildtype MEFs and human epithelial cells **(Figure 6)**. These results further express the value of mechanistic characterization and temporal dissection of gene expression shifts in driving rational drug discovery.

**Figure 2.**
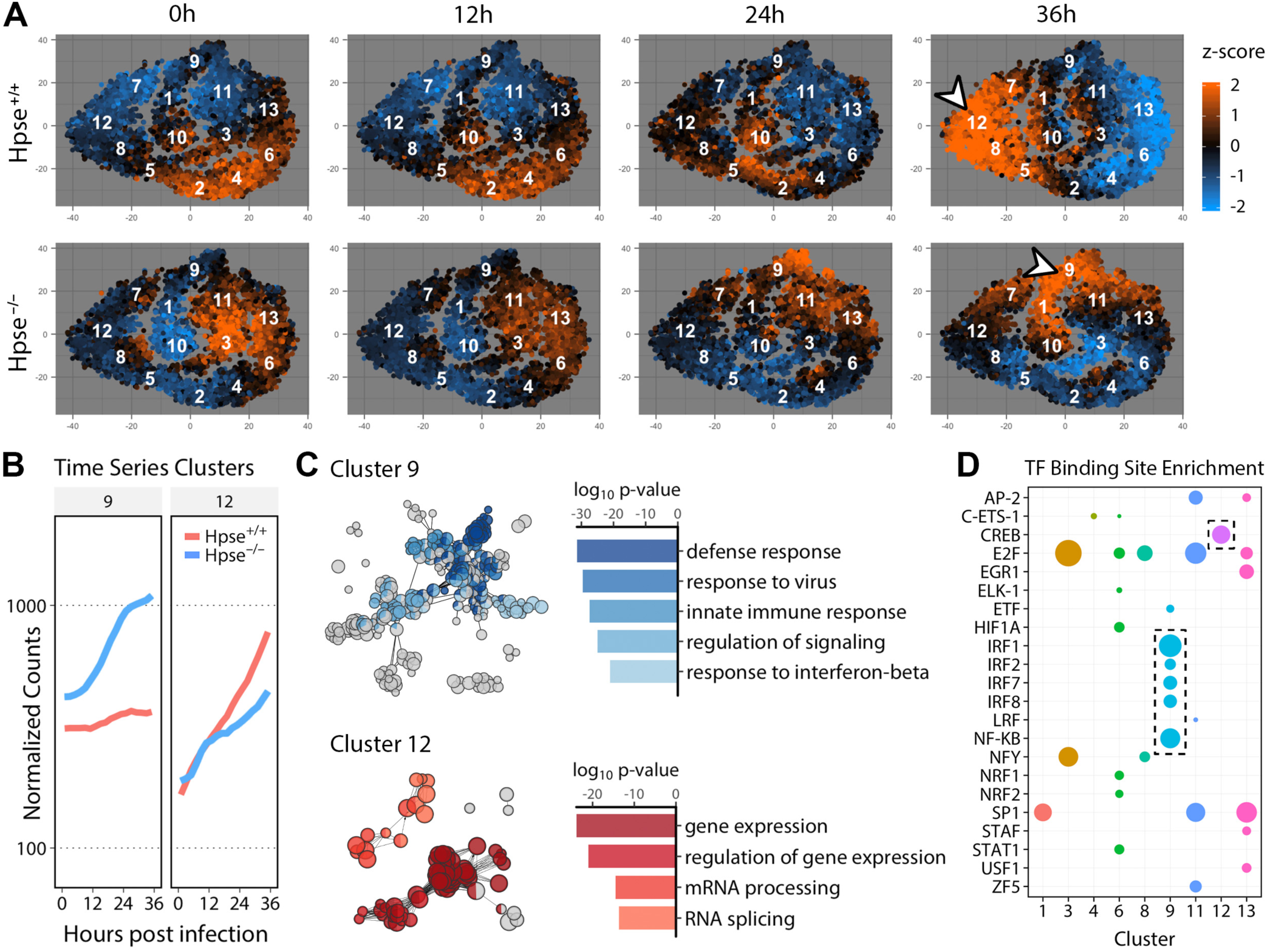
Temporal viromics catalogs shifts in transcriptional landscape and reveals functional clusters of genes dependent on HPSE. **(A)** t-stochastic neighbor embedding (t-SNE) projection depicting relative expression of differentially expressed genes, with clustering based on expression patterns over infection timecourse. Centroids of each of 13 clusters are labelled with respective identifier. Arrowheads indicate clusters further dissected below. **(B)** Median gene expression of selected clusters displayed over time. **(C)** Representation of significantly enriched gene ontology (GO) terms performed with ClueGO. Node sizes relative to GO term enrichment, with functionally related terms grouped by color and indicated by corresponding bar plot. **(D)** Significant transcription factor binding site enrichment analysis represented by cluster, with bubble sizes relative to enrichment score. See also Figures S1 and S3.

### HPSE restricts direct antimicrobial activity of type I interferon system

Given the amplified induction of defense and immune-related genes observed in our unbiased analysis of Hpse-deficient cells, we sought to mechanistically define the apparent link between HPSE and innate immune signaling. Subsetting both transcriptomics and proteomics datasets based on the curated “Hallmark Interferon Alpha Response” gene set (MSigDB) shows the striking upregulation of interferon response genes (ISGs) in the absence of HPSE **(Figures 3A and 3B)**. Interferons (IFN) are a highly conserved family of cytokines, with type I IFN (α and β) secreted particularly in response to viral infection and known to exert various antimicrobial functions (17). Using ISG15 as an indicator of signaling activity downstream of type I IFN, Hpse-KO cells are observed to be intrinsically more sensitive to these conserved cytokines. In the absence of HPSE, cells are responsive to levels of purified IFN-β orders of magnitude lower than wildtype cells, with active signaling occurring even without any external stimulus **(Figure 3C)**. Interestingly, with a similar size and structure to that of ubiquitin, ISG15 is known to exert antimicrobial activity through direct protein ligation, with downstream consequences including proteasomal degradation and regulation of signal transduction (18). Expression analysis of individual viral proteins by proteomics analysis showed that Hpse-KO cells exhibit a particularly large defect in production of the immediate early protein ICP0, which is known to be essential for HSV-1 replication and assembly **(Figures 3D-3F, S5)** (19, 20). Blocking of the proteasome with MG132 for the last 4 hours of infection somewhat restores ICP0 (110 kDa) and also reveals increased levels of a 66 kDa degradation product previously observed by other investigators (21). Our findings here detail one mechanism of direct antimicrobial action occurring in the absence of HPSE: conjugation of ISG15 to ICP0 correlates with increased proteasomal degradation of this essential viral protein, thus stalling viral replication **(Figure 3G)**.

**Figure 3.**
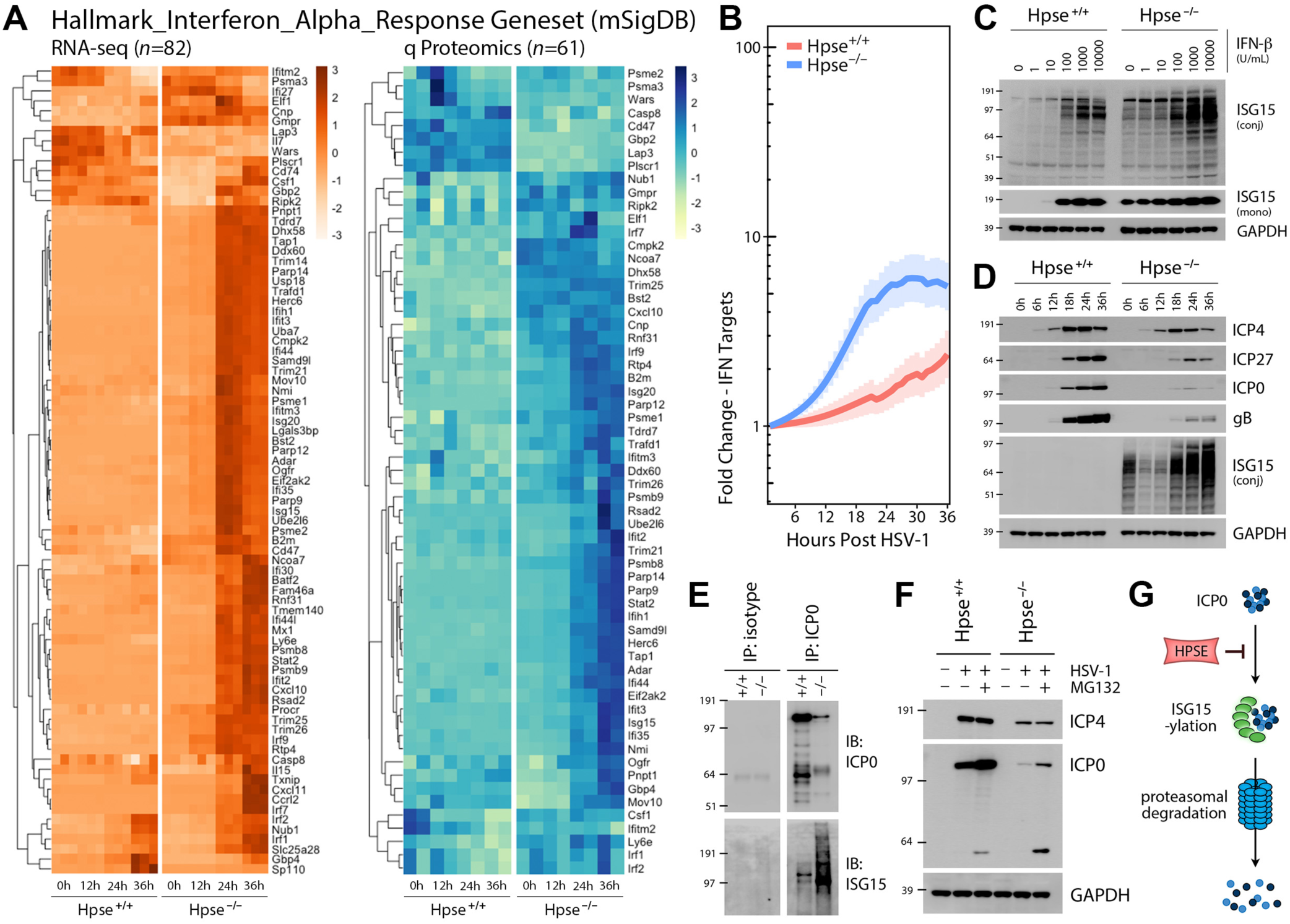
HPSE restricts type I interferon response and drives virus production through inhibition of degradation of immediate early protein ICPO. **(A)** Transcriptomic and proteomic datasets filtered based on Hallmark_Interferon_Alpha_Response geneset (*n*=91 genes) obtained from mSigDb. **(B)** Transcript expression of Hallmark_Interferon_Alpha_Response subset displayed as fold change over baseline, with bolded lines signifying mean expression and shaded areas indicating standard error. Splines generated using MetaLonDA. **(C)** Increased type I interferon sensitivity in Hpse ^-/-^ cells at baseline and with interferon (IFN)-β administration for 18 h, exemplified by expression of interferon stimulated gene (ISG)15 monomer and protein-conjugated forms. **(D)** Representative western blot analysis of cell lysates from wildtype and Hpse ^-/-^ MEFs indicating selective defect in viral protein production in HPSE deficiency. gB is a late (γ) gene, while ICP4, ICP27, and ICP0 are immediate early (α) genes. **(E)** Western blot analysis of cell lysate, isotype immunopurification (IP), ICP0 IP of wildtype and Hpse ^-/-^ cells after 24 h HSV-1 infection, with 10 μM MG132 added for the final 4 h. **(F)** Immediate early viral protein expression in the presence of proteasome inhibitor MG132 (10 μM for last 4 h of 24 h infection) shown by western blot analysis of cell lysates. **(G)** Proposed model depicting ability of HPSE to interfere with ISG15 conjugation and proteasomal degradation of viral ICP0.

### Deletion of HPSE protects from cellular infiltration and associated inflammation

To appreciate a broader impact of these findings, we evaluated infection in Hpse-KO mice (22), which showed a significant decrease in virus production by plaque assay after corneal HSV-1 infection **(Figure 4A)**. Despite the more modest reduction in virus titers compared to observations *in vitro*, HSV-1 infected corneas displayed a striking ablation of the typical inflammatory response observed in wildtype mice, evidenced by loss of CD45^+^ Gr-1^+^ neutrophil and monocyte infiltration at 7 and 14 dpi **(Figures 4B-4E)**. Infected Hpse-KO mice also showed a significant increase in corneal IFN-β mRNA, in line with our *in vitro* findings **(Figure 4F)**. Furthermore, ocular application of monoclonal antibodies against interferon-α receptor (α-IFNAR) resulted in a significant restoration of virus production and infiltrative phenotypes, including increased cellularity of draining lymph nodes and marked corneal opacity **(Figures 4G-4J)**. These results may suggest that resident corneal cells lacking HPSE are intrinsically more effective in neutralizing the virus rather than inciting the nonspecific granulocytic infiltration known to be pathogenic in herpes keratitis. Collectively, these findings demonstrate that HPSE restriction of the type I IFN system is a potent immunomodulatory mechanism, and upregulation of HPSE may be a common microbial strategy for evasion of innate immune responses.

**Figure 4.**
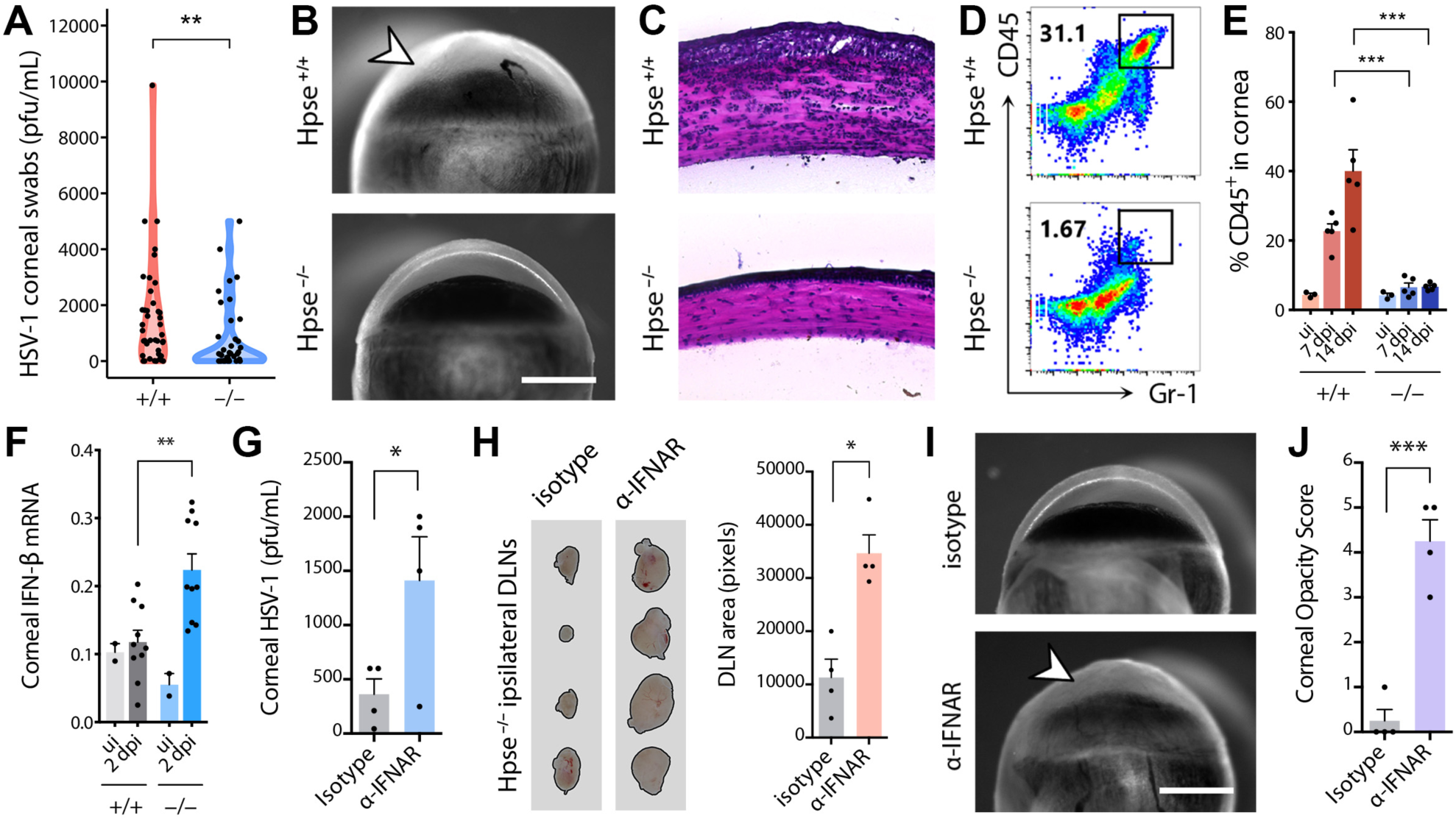
Deletion of HPSE protects from cellular infiltration and associated inflammation in a murine model of corneal infection. **(A)** HSV-1 titers 2 d following corneal infection of Hpse ^+/+^ and Hpse ^-/-^ mice (*n*=44). **(B-D)** Markedly decreased corneal inflammation and infiltration (arrowhead) in Hpse-deficient mice 14 d following HSV-1 infection observed by gross imaging (B), hematoxylin-eosin histology (C), and flow cytometry of dissociated corneal tissues (D). Scale bar, 1 mm. **(E)** Quantification of total leukocytes (CD45^+^) present in corneal tissue at indicated day post infection (dpi) as observed by flow cytometry. **(F)** Corneal IFN-β mRNA copy number relative to β-actin measured at 2 dpi (*n*=10). Uninfected (UI) samples constituted 2 pools of 5 mouse eyes each. **(G-J)** Partial restoration of viral titers and cellular infiltration in HPSE-deficient mice after topical application of α-IFNAR monoclonal antibody, observed by ocular wash titers (G), gross analysis of ipsilateral draining lymph nodes (DLN) (H) and gross analysis (I) and scoring (J) of corneal opacity (*n*=4). Scale bar, 1 mm. *p<0.05, **p<0.01, ***p<0.001.

### Necroptosis is an interferon-induced stress response limited by HPSE

Another initial observation from our studies was that Hpse-deficient cells appear to possess the intrinsic ability to contain infection before considerable viral spread can occur. This finding suggests that in the absence of HPSE, infected cells undergo some form of programmed cell death to shut down virus production. Indeed, Hpse-KO cells are more prone to virus-induced cell death, and possibly exhibit increased levels of basal cell death, measured as increased PI uptake by flow cytometry **(Figure 5A)**. Although effective viral production is markedly hindered in the absence of HPSE, small clusters of rounded and dead cells similar to plaques are frequently observed after infection **(Figure 5B)**. Interestingly, recent studies have suggested that necroptosis, or programmed necrosis, is a major mechanism of virus-induced cell death, which may provide some level of protection to the infected host (23-25). Moreover, type I IFNs are known drivers of necroptosis in response to various cellular stresses including viral infection (17). Western blot analysis of an infection time course showed a pronounced increased abundance of a key protein required for induction of necroptosis, receptor-interacting serine/threonine-protein kinase 3 (RIPK3), in the absence of HPSE **(Figure 5C)**. Increased abundance of mixed lineage kinase domain-like pseudokinase (MLKL) was also observed, while no apparent difference in other key proteins including RIPK1 and Caspase 8 was noted. Using inhibitors of the type I IFN system (α-IFNAR) and necroptosis (Nec-1) in combination, infection in Hpse-KO cells is nearly restored to wildtype levels **(Figures 5D-5F)**. Evaluation of inhibitors of apoptosis (ZVAD) and necroptosis (Nec-1) shows that HPSE preferentially limits necroptosis to promote cell survival and thus virus production. As many viruses including HSV-1 are known to block cell death to promote viral production and spread, clearance of infected cells through HPSE inhibition may hold great therapeutic potential.

**Figure 5.**
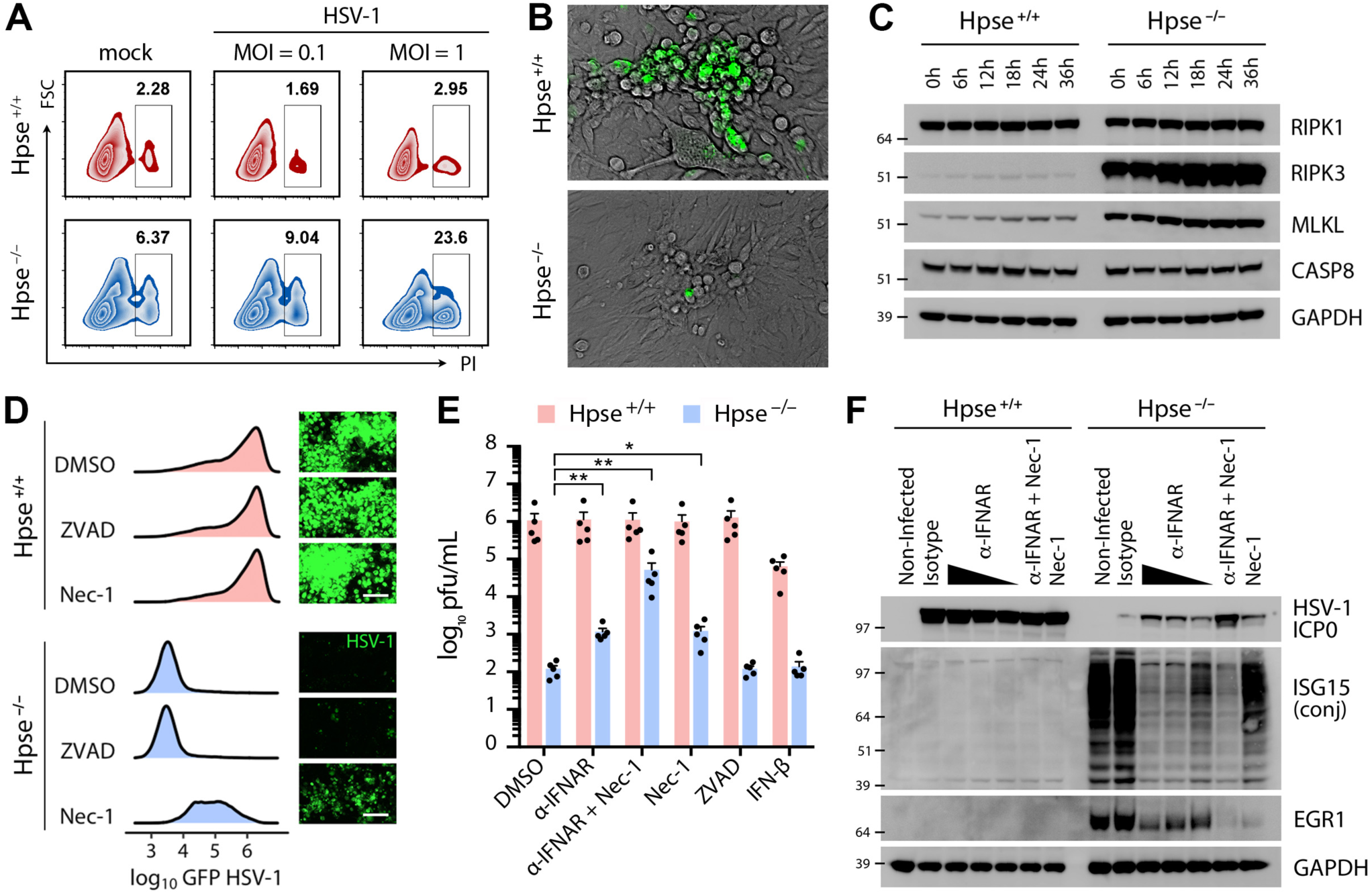
Attenuation of interferon response and necroptotic cell death restores infection in the absence of HPSE. **(A)** Measurement of cell death by flow cytometry detection of propidium iodide (PI) cellular uptake after 24 h HSV-1 or mock infection. **(B)** Immunofluorescence microscopy of cells infected with GFP-HSV-1, images captured at 24 hpi. Despite profound abrogation of virus production in absence of HPSE, multiple clusters of rounded and detached cells resembling plaques are observed after infection. **(C)** Western blot analysis of key proteins involved in induction of necroptosis at indicated times post infection. **(D)** Representative flow cytometry quantification and micrographs of Hpse ^+/+^ and Hpse ^-/-^ cells after infection with GFP-HSV-1 for 48 h, incubated with inhibitors of apoptosis (ZVAD) or necroptosis (Nec-1) as indicated. Scale bar, 100 μm. **(E)** Restoration of virus production in Hpse ^-/-^ cells with blocking of interferon alpha receptor (α-IFNAR) and necroptosis (Nec-1) showing a synergistic effect (*n*=5). **(F)** Inverse relationship between viral infected cell protein (ICP0) and ISG15 expression demonstrated by western blot, with near complete rescue of virus production observed upon α-IFNAR and Nec-1 treatment in Hpse ^-/-^ cells. *p<0.05, **p<0.01.

### Bioinformatics-guided analysis of transcription factor activation in HSV-1 infection identifies potent antiviral compounds

While gaining an understanding of the major antiviral pathways active in Hpse-KO cells, we remained interested in targeting proviral networks activated upon infection of wildtype cells. Analysis of divergent responses between these two cell types provided the initial clues that cAMP response element-binding protein (CREB) and β-catenin are drivers of proviral signaling. CREB was identified as a key driver of cluster 12 expression through transcription factor binding site analysis of RNA-seq data **(Figure 2)**, while β-catenin was detected as a major hub gene from proteomics analysis of differentially expressed proteins. In light of these results and the above findings that HPSE inhibits functional interferon signaling, we pursued further experiments under the following rationale **(Figure 6A)**. Years of published literature informed us that interferon regulatory factors, including IRF3, IRF7 and IRF9, bind to CREB-binding protein (CBP) and histone acetyltransferase p300 to drive optimal transcription of type I IFN and downstream IFN-stimulated genes (26, 27). Thus, these IRFs in effect compete with CREB and other transcription factors for CBP/p300 occupancy and binding to respective cAMP response elements (CRE) or interferon-sensitive response elements (ISRE). Additionally, we focused some attention to EGR1, a transcription factor shown by multiple studies to drive HPSE upregulation (28, 29), and act as a supporter of IFN signaling (30, 31). Interestingly, various studies have also drawn connections of EGR1 with necroptosis, CREB signaling and viral infection (32-35).

**Figure 6.**
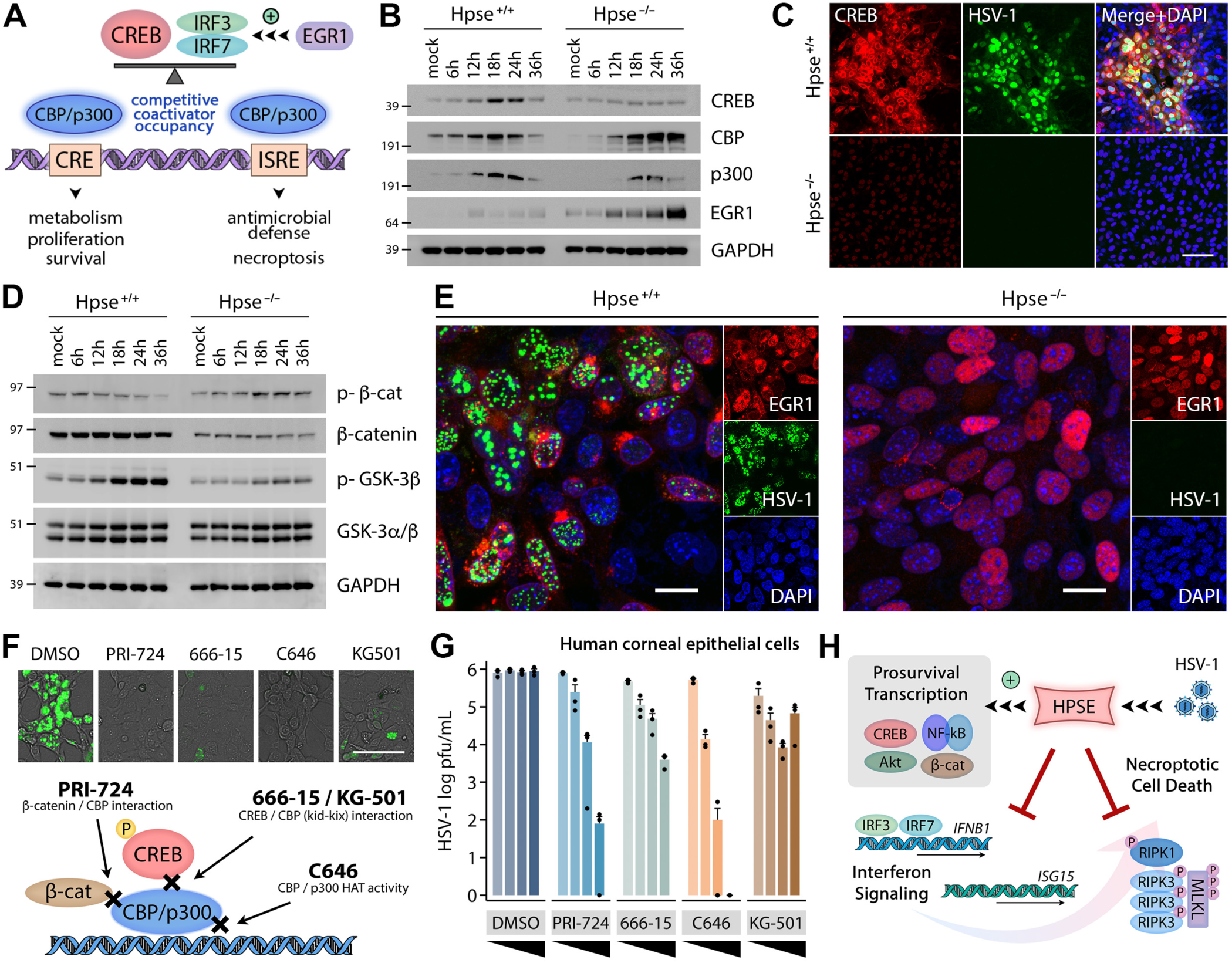
Bioinformatics-guided analysis of transcription factor activation in viral infection identifies potent antiviral compounds. **(A)** CREB and IRFs were identified by previous temporal viromics analysis as major regulators of prosurvival and defense response induction upon infection, respectively. cAMP response element binding (CREB) and interferon regulator factors (IRFs) are known to compete for occupancy of their common enhanceosome complex involving the transcriptional coactivators CBP/p300. Once respective complexes are formed, optimal DNA binding and transcription is achieved at cAMP response element (CRE) or interferon stimulated response element (ISRE) sites. Interestingly, the transciption factor early growth response 1 (EGR1) is reported to be both a supporter and a target of interferon signaling, and is also known to upregulate HPSE transcription. **(B)** Representative western blot analysis of CREB signaling induction with infection of wildtype and Hpse-KO cells. **(C)** Confocal immunofluorescence microscopy of wildtype and Hpse-KO cells showing CREB upregulation in infected wildtype cells. Scale bar, 100 μm. **(D)** Representative western blot analysis of β-catenin signaling induction with infection of wildtype and Hpse-KO cells. **(E)** Confocal immunofluorescence microscopy of wildtype and Hpse-KO cells showing EGR1 cellular localization and GFP-HSV-1. Scale bar, 20 μm. **(F)** *Top*, Representative immunofluorescence micrographs of human corneal epithelial cells infected with GFP-HSV-1 and then incubated with specified inhibitors at 2 h post GFP-HSV-1 infection, images captured at 24 hpi. Scale bar, 100 μm. *Bottom*, Schematic depicting known mechanisms of action of selected inhibitors. **(G)** Viral titers obtained after incubation of specified inhibitors at concentrations 12.5, 25, 50 and 100 μM. (666-15 was used at concentrations of 5, 10, 15, and 20 μM due to unacceptable toxicity at higher levels). **(H)** Model of HPSE function at the interface of innate defense responses and cell survival. Infection and other cellular insults trigger activation and nuclear translocation of multiple prosurvival factors, including CREB, Akt, NF-κB and β-catenin. Previous work and this study show that HPSE modulates nuclear trafficking of these TFs, which drive cellular proliferation, microbial replication and carcinogenesis. Here we show that HPSE inhibits type I interferon production and induction of necroptosis, in part via EGR1 which has been previously associated with each of these processes. These innate stress responses likely act to protect multicellular tissues from viral or cancerous spread by preventing uncontrolled cellular proliferation.

Indeed, western blot and immunofluorescence microscopy confirmed that activation of the CREB and β-catenin systems upon infection appears unique to wildtype MEFs, while Hpse-KO cells do not display these changes **(Figures 6B-6D)**. CREB is upregulated in wildtype cells starting at 12 hpi, while this response is absent in Hpse-KO. Likewise, phosphorylation of β-catenin at the S33/S37/T41 sites, indicative of inactive β-catenin (36), is decreased across infection timepoints, while this phosphorylation is increased in Hpse-KO. A similar trend is observed for p-GSK3β, which when phosphorylated at the S9 site is known to result in β-catenin activation through double inhibition (36). Given these supportive findings, we opted to treat cells with inhibitors targeting the CREB or β-catenin systems by therapeutic application at various concentrations at 2 hpi after viral entry had occurred. The identities and mechanisms of these compounds are shown in **Figure 6F**. Initial testing was performed in wildtype MEFs; upon detection of antiviral efficacy we transitioned to human corneal epithelial (HCE) cells, a model cell line for HSV-1 infection. Images from GFP-HSV-1 and viral titers from HCE cell culture supernatants collected at 24 hpi demonstrate antiviral efficacy of these compounds **(Figures 6F and 6G)**. By merging our unbiased temporal analysis with molecular manipulation, we show that through activation of these two host factors with broad control over cellular physiology, HSV-1 generates an environment conducive to its own propagation and persistence.

## Discussion

With this multidisciplinary exploration of various cellular responses to stress, we demonstrate how HPSE serves as a key intersection between the detection and effector phases of signal transduction. Using a genome-wide analysis of HPSE-deficient cells, we aimed to define the factors and related pathways that make these cells naturally resistant to infection. Our results show that considerable differences in baseline gene expression exist in these cells compared to wildtype, possibly due to compensatory mechanisms that have emerged throughout embryonic development in the absence of HPSE. Viral infection triggers extensive remodeling of host cellular processes. By understanding the connections between these processes, we can infer their reliance on HPSE in the case of wildtype cells. The temporal viromics methods employed here reliably clustered differentially expressed genes and identified transcription factors regulating major shifts in cellular programs over time. This unique analysis thus draws an indirect connection between HPSE and multiple important drivers of cellular proliferation and defense. In agreement with our previous work, we show by transcription factor binding site analysis that IRFs and NF-κB are major drivers of the defense response gene cluster heavily upregulated with infection in Hpse-KO cells. We further dissect this relationship *in vitro* and *in vivo* by rescuing virus production in Hpse-KO with inhibition of the interferon and necroptosis systems. Of note, a majority of studies have used multiplicities of infection (MOI) of HSV-1 in the 1 to 100 range for induction of interferon signaling in MEFs. In this study, we focused on an MOI of 0.1, more commensurate to productive infection occurring *in vivo*. Our results highlight the increased sensitivity of Hpse-KO cells to induction of the type I interferon response upon low grade viral infection. Future studies will aim to more precisely define the nature of the first-order interactions between HPSE and effectors as these remain almost entirely unknown. Given the regulatory relationship that exists between EGR1 and HPSE (28, 29), we suspect that in the absence of HPSE expression, EGR1 upregulation upon infection remains unchecked, and that this uncontrolled EGR1 level results in hyperactivation of the type I IFN system. Further mechanistic studies will be required to dissect the antiviral mechanisms displayed by EGR1. Moving forward, high-throughput analyses with even greater temporal resolution will continue to unearth unique connections not previously imagined, yet the importance of molecular validation by conventional methods will remain. Additionally, these newly reported activities of HPSE can help explain the documented roles of HPSE in cancer progression. Numerous malignancies have been shown to increase expression of HPSE, understood to drive late stage metastatic potential (1, 2). Recent studies have also shown that the type I interferon system has control over cancer spread as depletion of IRF7 increases metastatic burden in mice and humans (37, 38).

We also identify two novel proviral transcriptional regulatory factors CREB and β-catenin that are activated in wildtype cells, and demonstrate that their pharmacological inhibition in human cells is a highly effective antiviral strategy resulting from this analysis. We show that through activation of these two factors with broad control over cellular physiology, the virus generates an environment conducive to its own success. CREB is a transcription factor well studied in the neuroscience field, but with little mention in studies of microbial pathogenesis. It is known to be activated by protein kinases upon second messenger signaling and bound by coactivators CREB-binding protein (CBP) and p300 for its binding to DNA sequences (39). CREB has also been documented to interact with various signaling molecules including signal transducer of and activator of transcription (STAT1) and NF-κB, indicating potential ramifications for generation of immune responses (40, 41). Likewise, β-catenin is a well-studied regulator of transcription and cell adhesion overexpressed in several cancers, which is part of a multi-protein destruction complex. Numerous protein interactions and regulators of β-catenin have been characterized, with one known transcriptional co-activator being CBP for DNA binding. The association between this prosurvival factor and infection remains understudied as only a small number of studies exist linking HSV-1 or other viral infection to β-catenin activation (42-44). In this study, we show that multiple commercially available compounds targeting CREB and β-catenin systems demonstrate antiviral efficacy against HSV-1. While some degree of cellular toxicity was noted at higher concentrations, all compounds tested displayed antiviral efficacy, with PRI-724 showing the most appealing profile of antiviral activity over toxicity. Interestingly, this compound is currently in clinical trials for hepatitis C virus hepatocellular carcinoma and other cancers (45), and along with the other compounds analyzed here may prove a promising antiviral candidate. Our findings showcase a success of mechanism-based rational drug design informed by multi-perspective analysis merging systems and molecular investigation.

## Materials and Methods

### Cell lines and virus strains

Human corneal epithelial (HCE) cell line (RCB1834 HCE-T) was obtained from Kozaburo Hayashi (National Eye Institute, Bethesda, MD) and was cultured in MEM (Life Technologies, Carlsbad, CA) with 10% fetal bovine serum (FBS) (Life Technologies) and 1% penicillin/streptomycin (Life Technologies). Confirmation of identity of HCE cell line was done by short tandem repeat analysis. Vero cell line for virus preparation and plaque assay was provided by Dr. Patricia G. Spear (Northwestern University, Chicago, IL) and cultured in DMEM (Life Technologies) with 10% FBS and 1% penicillin/streptomycin. Wildtype and heparanase-knockout mouse embryonic fibroblasts (WT and Hpse-KO MEFs) were provided by Dr. Israel Vlodavsky (Rappaport Institute, Haifa, Israel) (22). All cells were maintained in a Heracell VIOS 160i CO2 incubator (Thermo Scientific) and have been confirmed negative for mycoplasma contamination. Virus strains used in these studies, HSV-1 (KOS-WT) (46), GFP-HSV-1 (K26-GFP) (46), HSV-1 (gL86), McKrae and pseudorabies virus (PRV) (47) were provided by Dr. Patricia G. Spear (Northwestern University, Chicago, IL). Dual color ICP0_p_-GFP/gC_p_-RFP was a gift of Dr. Paul Kinchington (University of Pittsburgh). HSV-1 infections were performed with strain KOS at MOI=0.1 unless otherwise specified. All virus stocks were propagated in Vero cells and stored at −80°C.

### Antibodies and plasmids

The following antibodies were used for western blot studies: from Cell Signaling Technology, at dilution 1:1000 – CREB (9197), CBP (7389), p300 (86377), β-catenin (8480), phospho-β-catenin (9561), EGR1 (4154), RIPK1 (3493), RIPK3 (95702), MLKL (37705), CASP8 (4927); from Santa Cruz Biotechnology, at dilution 1:250 – GSK-3β (sc-7291), phospho-GSK-3β (sc-373800), ICP0 (sc-53070), ICP4 (sc-69809), ICP27 (sc-69806), ISG15 (sc-166755); from Proteintech, at dilution 1:1000 – GAPDH (10494); from Abcam, at dilution 1:10,000 – gB (ab6505). The following antibodies were used for immunofluorescence microscopy studies: from Cell Signaling Technology, at dilution 1:100 – CREB (9197), EGR1 (4154). HPSE expression constructs including Myc-GS3 plasmid were provided by Dr. Israel Vlodavsky (Rappaport Institute, Haifa, Israel) (48). Lipofectamine-2000 transfection reagent (Life Technologies, 11668019) was used for all *in vitro* overexpression experiments, according to the manufacturer’s specifications.

### Chemical reagents

Anti-mouse interferon alpha receptor 1 (α-IFNAR) was purchased from Leinco (St. Louis, MO) and was used at concentrations ranging from 0.1 μg/mL to 10 μg/mL, as indicated (I-401, clone MAR1-5A3). Mouse isotype control antibody (mouse IgG1, Cell Signaling 5415) was used as a negative control for this antibody where applicable. Necrostatin-1 (Nec-1) was used to inhibit necroptosis and was purchased from Selleckchem (Houston, TX) and used at 50 μM, unless otherwise specified (S8037). Pan-caspase inhibitor Z-VAD-FMK was used at 10 μM to block apoptosis and was purchased from Selleckchem (S7023). MG132 was used at 10 μM to inhibit proteasomal degradation of proteins and was purchased from Selleckchem (S2619). Pharmacological inhibitors of the CREB/CBP/β-catenin system were purchased from Selleckchem (PRI-724, S8262; KG-501, S8409; C646, S7152) and Tocris Biosciences (666-15, 5661). Purified mouse interferon-β was purchased from PBL Assay Science (Piscataway, NJ) (12405-1).

### Murine ocular infection model

All animal care and procedures were performed in accordance with the institutional and NIH guidelines, and approved by the Animal Care Committee at University of Illinois at Chicago (ACC protocol 17-077). 6-10-week-old male and female mice on C57/B6 background (wildtype or Hpse-deficient) were used for all experiments. Anaesthetized mouse corneas were scarified in a 3×3 grid using a 30-gauge needle, and infected as previously described (11). All images of the corneal surface were acquired with SteREO Discovery.V20 stereoscope (Zeiss, Germany). For IFNAR blockade experiment, 5 μL α-IFNAR monoclonal antibody (Leinco I-401) was applied topically to mice corneas 24 h prior to infection and then daily for 3 days post infection. Ocular infection was performed as described above and mice tissues were analyzed at 45 dpi.

### Western blot

Cellular proteins were extracted using the following lysis buffer: 150 mM NaCl, 50 mM Tris-HCl pH 7.4, 10% glycerol, 1% NP-40, 10 mM sodium fluoride, 1 mM sodium orthovanadate, 10 mM N-ethylmaleimide, and Halt Protease Inhibitor Cocktail (Thermo Scientific). Lysis was performed on ice with agitation for 30 min, followed by 30 min centrifugation at 13,000 rpm. Clarified lysates were then denatured at 95°C for 8 min in the presence of 4X LDS sample loading buffer (Life Technologies) and 5% beta-mercaptoethanol (Bio-Rad, Hercules, CA) and separated by SDS-PAGE with NuPAGE 4-12% Bis-Tris 1.5 mm 15-well gels (Thermo Scientific). Proteins were then transferred to nitrocellulose using iBlot2 system (Thermo Scientific) and membranes were blocked in 5% milk/TBS-T for 1 h, followed by incubation with primary antibody overnight. After washes and incubation with respective horseradish peroxidase-conjugated secondary antibodies (Jackson ImmunoResearch Peroxidase AffiniPure goat anti-mouse IgG (H+L), 115-035-146 at 1:10,000 or Peroxidase AffiniPure goat anti-rabbit IgG (H+L), 111-035-003 at 1:20,000) for 1 h, protein bands were visualized using SuperSignal West Femto substrate (Thermo Scientific) with Image-Quant LAS 4000 biomolecular imager (GE Healthcare Life Sciences, Pittsburgh, PA).

### Quantitative polymerase chain reaction

RNA was extracted from cultured cells using TRIzol (Thermo Scientific, 15596018), following the manufacturer’s protocol, and complementary DNA was produced using High Capacity cDNA Reverse Transcription kit (Life Technologies). Where applicable, corneal tissues were extracted from mice and incubated in 50 μL of 2 mg/mL collagenase D (Sigma C0130) in PBS for 1 h at 37°C. Tissues were then triturated with a pipet tip, resuspended in TRIzol and extraction of RNA and cDNA were performed as above. Real-time quantitative polymerase chain reaction (qPCR) was performed using Fast SYBR Green Master Mix (Life Technologies) on QuantStudio 7 Flex system (Life Technologies). To quantify viral genomes, infected cell pellets were resuspended in 500 μL virion buffer: 8 mg/mL Tris-HCl, 1% SDS, 10 mM EDTA supplemented with 1 μL proteinase K (Thermo Scientific EO0491) and incubated at 55°C for 16 h. Viral DNA was then extracted by adding 500 μL of Ultrapure phenol chloroform isoamyl alcohol mix (Thermo Scientific 15593-031), according to the manufacturer’s specifications. The gD-specific primers listed below were then used to quantify HSV-1 genomes.

The following mouse-specific primers were used in this study:

IFN-β Fwd 5’-TGTCCTCAACTGCTCTCCAC-3’, Rev 5’-CATCCAGGCGTAGCTGTTGT-3’

ISG15 Fwd 5’-AGCAATGGCCTGGGACCTAAAG-3’, Rev 5’-CCGGCACACCAATCTTCTGG-3’

β-actin Fwd 5’-GACGGCCAGGTCATCACTATTG-3’, Rev 5’-AGGAAGGCTGGAAAAGAGCC-3’

The following HSV-1-specific primers were used in this study:

gD Fwd 5’-GTGTGACACTATCGTCCATAC-3’, Rev 5’-ATGACCGAACAACTCCCTAAC-3’

ICP0 Fwd 5’-ACAGACCCCCAACACCTACA-3’, Rev 5’-GGGCGTGTCTCTGTGTATGA-3’

### Plaque assay

Cell monolayers were inoculated for 2 h in Opti-MEM (Thermo Scientific) with incubation at 37°C, 5% CO_2_, after which viral suspension was aspirated and replaced with complete media for remaining infection time course. To quantify extracellular virus, culture supernatants were collected, centrifuged at 13,000 rpm for 1 min, serially diluted in Opti-MEM, and overlaid on confluent monolayers of Vero cells in 24-well plates. After 2 h incubation, plaque inocula were aspirated, and cells were incubated with complete DMEM containing 0.5% methylcellulose (Fisher Scientific) for 48 to 72 h. Medium was then aspirated, cells were fixed with 100% methanol and finally stained with crystal violet solution to visualize plaques.

### Viral β-galactosidase entry assay

Cell monolayers were inoculated with recombinant HSV-1 strain gL86, in which a portion of the gL gene was replaced with the lacZ gene encoding β-galactosidase. Only upon successful entry and expression of the viral genome is the β-galactosidase enzyme produced, and this virus is only capable of one round of viral replication; newly produced virions will not be capable of downstream infection. At 6 h post infection, cells were washed once with PBS and incubated at 37°C for 1 h with ONPG solution (3 mg/mL (Fisher Scientific, 34055) + 0.05% NP-40) and colorimetric substrate was detected at 410 nm using a microplate reader (Tecan GENious Pro, Mannedorf, Switzerland).

### Immunofluorescence microscopy

HCE cells or wildtype and Hpse-KO MEFs were cultured in glass bottom dishes (MatTek Corporation, Ashland, MA) or 8-well μ-slides (iBidi, Madison, WI). Cells were fixed in 4% paraformaldehyde for 10 min and permeabilized with 0.1% Triton-X for 10 min at room temperature for intracellular labeling. This was followed by incubation with primary antibody for 1 h at room temperature. When a secondary antibody was needed, cells were incubated with respective FITC- or Alexa Fluor 647-conjugated secondary antibody (Sigma-Aldrich F9137 or Thermo Scientific A21244) at a dilution of 1:100 for 1 h at room temperature. NucBlue Live ReadyProbes Hoescht stain (Thermo Scientific R37605) was included with secondary antibody stains when applicable, according to manufacturer’s specifications. Samples were examined under LSM 710 confocal microscope (Zeiss) using a 63X oil immersion objective. For cell surface staining, cells were incubated with respective antibodies prior to fixation with paraformaldehyde for imaging. Fluorescence intensity of images was calculated using ZEN software.

### Live cell imaging

Wildtype and Hpse-KO MEFs were cultured in 24-well plates and infected with HSV-1 dual color fluorescent virus expressing GFP driven by ICP0 promoter and RFP driven by gC promoter. At 2 hpi, inoculation media was replaced with complete DMEM, and cells were placed in the incubation chamber of Zeiss spinning disk live-cell imaging system, which maintains conditions of 37°C and 5% CO_2_. Images were captured on RFP, GFP, and brightfield at an interval of 30 min for 36 h, and analyzed with ZEN software.

### Immunopurification

Immunopurification (IP) of proteins was performed using HCE cells cultured in 15 cm dishes. Cells were collected after specified times of infection and/or treatment, washed with PBS, scraped in cold PBS on ice and transferred to conical tubes. Cells were centrifuged at 1200 rpm for 5 min, then cellular proteins were extracted with lysis buffer: 150 mM NaCl, 50 mM Tris-HCl pH 7.4, 10% glycerol, 1% NP-40, 10 mM sodium fluoride, 1 mM sodium orthovanadate, 10 mM N-ethylmaleimide, and Halt Protease Inhibitor Cocktail (Thermo Scientific). After 30 min of lysis with agitation at 4°C, lysates were centrifuged at 13,000 rpm for 30 min to pellet insoluble cellular debris. IP antibody, ICP0 (Santa Cruz, sc-53070) or isotype control (mouse IgG1, Cell Signaling 5415) were then added to clarified lysates and rotated for 16 h at 4°C. Protein A/G Dynabeads (Thermo Scientific, 88802) were added and samples were rotated for 1 h at 4°C. Four washes with magnetic separation were performed with lysis buffer. Beads were finally resuspended in LDS buffer with 5% beta-mercaptoethanol, denatured at 95°C for 8 min, and SDS-PAGE was performed.

### Flow cytometry

Corneas were extracted from mice after euthanasia and treated with 2 mg/mL collagenase D (Sigma C0130) for 1 h at 37°C and triturated with a pipet tip. Cell suspensions were filtered through a 70 μm mesh, resuspended in FACS buffer (PBS + 5% FBS), and counted by hemocytometer. 10^6^ cells from each sample were aliquoted into U-bottom 96-well plates for subsequent staining. F_C_-receptors were blocked using TruStain fcX (101319, Biolegend, San Diego, CA), and the following fluorophore-conjugated primary antibodies from Biolegend were used for cell surface staining: FITC anti-mouse CD45 (103107) and APC anti-mouse Ly-6G/Ly-6C (Gr-1) (108411). Cells were immunolabeled, washed, and analyzed with Accuri C6 Plus flow cytometer (BD Biosciences). For flow cytometric quantification of cell death by PI uptake, cells were either infected with HSV-1 KOS (MOI=0.1, MOI=1) or mock treated for 24 h in medium containing PI. At the termination of cellular incubations, cells were collected on ice, washed twice with FACS buffer and analyzed with Accuri C6 Plus flow cytometer. BD Accuri C6 Plus software and Treestar FlowJo v10.0.7 were used for all flow cytometry data analysis.

### Mouse tissue sectioning and staining

For hematoxylin and eosin staining, mice whole eyes were extracted after euthanasia, washed in PBS, embedded in Tissue-Tek OCT compound (Sakura Finetek), frozen on dry ice and stored at −80°C. 10 μm sections were cut and adhered to Superfrost Plus glass slides (Fisher Scientific) using Cryostar NX50 microtome (Thermo Scientific), air dried at room temperature for 10 min and then fixed in ice-cold acetone for 5 min. Slides were then incubated in the following series: 2 min running water, 1 min Mayer’s hemalum solution (Merck HX73030749), 2 min running water, 2 min 70% ethanol, 1 min 100% ethanol, 1 min eosin Y solution with phloxine (Sigma HT110316), 2 min 70% ethanol, 1 min 100% ethanol, 1 min xylene, and finally mounted and cover slipped with permount. Slides were then imaged using Zeiss Axioskop 2 Plus microscope.

### RNA sequencing

Wildtype and Hpse-KO MEFs were cultured in biological triplicates in 10 cm dishes with HSV-1 infection using strain KOS (MOI=0.1) at timepoints of 0, 12, 24 and 36 hpi for a total of 24 samples. At specified timepoints, cells were washed once in PBS, and collected by scraping on ice. 10^6^ cells from each dish were used for RNA-seq workflow, and remaining 4×10^6^ cells were used for qProteomics; the two analyses were performed on the same sets of cells. Cell pellets were suspended in RNAlater (Thermo Scientific, AM7020) and stored at −80°C until further processing. RNeasy Mini kit (Qiagen, 74104) was used to extract RNA, and DNA libraries for sequencing were constructed using Nextera XT DNA Library Prep kit (FC-131-1024, Illumina, San Diego, CA). The 24 prepared libraries were pooled and submitted to Michigan State University Genomics Facility, where they were quality checked using Qubit dsDNA HS, Agilent Bioanalyzer DNA 1000 and Kapa Illumina Library Quantification qPCR assays. The pool of libraries was loaded onto one lane of an Illumina HiSeq 4000 flow cell and sequencing was performed in a 2 × 150 bp paired end format using HiSeq 4000 SBS reagents (Illumina) and approximately 312 million acceptable quality reads were achieved. Base calling was done by Illumina Real Time Analysis v2.7.7 and output was demultiplexed and converted to FastQ format with Illumina Bcl2fastq v2.19.1. Assembly and alignment to Mus musculus (GRCm38.p6) and HSV-1 (JQ673480.1) genomes was performed with CLC Genomics Workbench. Subsequent analyses were performed in *R* v3.5. Differential expression analysis and normalization was performed with DESeq2, with design = ∼ Genotype + Time.point + Genotype:Time.point and reduced model = ∼ Genotype + Time.point, using likelihood relatedness test. Clustering of significantly differentially expressed genes (alpha = 0.01) was performed with Rtsne (15) and APcluster (16) based on temporal expression pattern. Transcription factor binding site enrichment analysis was performed using the PASTAA algorithm (49). Time series splines were generated with MetaLonDA (50). Gene ontology analysis was performed with ClueGO v2.5.2 (51) within Cytoscape v3.6.1 and ClusterProfiler v3.10.0 (52) in *R*.

### Mass Spectrometry-Based Proteomic Sample Preparation

Proteomic samples were processed as previously described with minor alterations (53). Cell pellets from HSV-1 infection time course were resuspended in 500 μL lysis buffer (75 mM NaCl, 3% SDS, 1 mM NaF, 1 mM β-glycerophosphate, 1 mM sodium orthovanadate, 10mM sodium pyrophosphate, 1 mM PMSF, 50 mM HEPES pH 8.5, and 1X Roche Complete EDTA-free protease inhibitor cocktail, followed by addition of 500 μL solution of 8 M urea + 50 mM HEPES pH 8.5, and solubilized by probe sonication. Samples were next subjected to reduction of disulfide bonds in 5 mM DTT for 30 min at 56°C. Free disulfide bonds were then alkylated for 20 min in the dark via the addition of iodoacetamide to a final concentration of 15 mM, followed by addition of the original volume of DTT and 15 min incubation in the dark. One quarter volume of trichloroacetic acid was then added to precipitate proteins. Samples were incubated on ice for 10 min, then centrifuged for 5 min at 18,000 x g at 4°C. The supernatant was removed and protein was washed twice via addition of ice-cold acetone, followed by centrifugation at 18,000 x g for 2 min between each wash. Pellets were dried and resuspended in a solution of 1 M urea + 50 mM HEPES pH 8.5 for digestion, performed in two steps. First, samples were incubated overnight at room temperature on a shaker in presence of 3 μg LysC Endopeptidase (VWR). Next, samples were incubated for 6 h at 37°C with 3 μg Sequencing Grade Modified Trypsin (Promega). Digestion was terminated via addition of 20 μL trifluoroacetic acid (TFA). Samples were subjected to centrifugation at 18,000 x g for 5 min to pellet insoluble debris, and supernatants were desalted using C18 resin columns (Waters) and samples were dried under vacuum. Extracted peptides were resuspended in a solution of 50% acetonitrile + 5% formic acid (FA) and quantified using a Quantitative Colorimetric Peptide Assay (Pierce), as recommended by the manufacturer. 50 μg of each sample was separated for subsequent sample preparation. A pooled “bridge channel” was made containing equal amounts from each sample, and two 50 μg aliquots were separated for inclusion in subsequent sample preparation steps. 50 μg aliquots were dried under vacuum. 50 μg aliquots were next labeled using tandem mass tags (TMT) (Thermo Scientific, 90309). Dried peptides were resuspended in a solution of 30% dry acetonitrile (ACN) + 200 mM HEPES pH 8.5. Label assignment was performed so that each biological replicate was represented within each 10-plex, and such that no two replicates were assigned to the same label, except for the bridge channel internal standards. 8 μL of 20 μg/μL TMT solution was added to the appropriate assigned sample, and the labeling reaction was allowed to proceed for 1 h at room temperature. Reaction quenching was performed via addition of 9 μL 5% hydroxylamine at room temperature for 15 min. 50 μL 1% TFA was added to each sample, and samples within each 10-plex experiment were multiplexed, desalted and vacuum dried as described above. 10-plexes were fractionated using reverse phase high pH liquid chromatography. Samples were resuspended in 110 μL 5% ACN + 5% FA. Fractionation was performed on an Ultimate 3000 HPLC with 4.6 mm x 250 mm C18 column. Samples were separated on a 1 h solvent gradient ranging from 5% to 90% ACN + 10 mM ammonium bicarbonate. Fractions were pooled as previously described and dried under vacuum (54). Alternating pooled samples were used for subsequent analysis, except for the tenth pooled fraction, where its complement was used.

### Mass Spectrometry-Based Proteomic Analysis

Dried fractions were resuspended in 8 μL 5% ACN + 5% FA and analyzed on an Orbitrap Fusion with in-line Easy Nano-LC 1000 (Thermo Scientific). Fractions were run as 3 h gradients progressing from 3% ACN + 0.125% FA to 100% ACN + 0.125% FA. Fractions were loaded onto an in-house packed column of 0.5 cm C4 resin with 5 μm diameter, 0.5 cm C18 resin with 3 μm diameter, and 29 cm of C18 resin with 1.8 μm diameter. The column measured 30 cm in overall length, with inner diameter of 100 μm and outer diameter of 360 μm. Electrospray ionization was achieved at the source through application of 2,000 V of electricity though a T-connector at the junction between the source, waste, and column capillaries. Spectra were acquired at the MS1 level in data-dependent more with scan range in the Orbitrap between 500 and 1200 m/z and resolution of 60,000. Automatic gain control (AGC) was set to 2 × 10^5^ with the maximum ion inject time set to 100 ms. The Top N setting was used for subsequent fragment ion analysis, with N=10. MS2 data collection was performed using the decision tree option, wherein ions with 2 charges were analyzed between 600-1200 m/z. Ions with 3-4 charges were analyzed between 500-1200 m/z. The lower ion intensity threshold was set to 5 × 10^4^, and selected ions were isolated in the quadrupole at 0.5 Th. Fragmentation was performed using Collision-Induced Dissociation, and ion detection and centroiding occurred in the linear ion trap with AGC rapid scan rate of 1 × 10^4^. MS3 data was generated using synchronous precursor selection (55). No more than 10 precursor ions were concurrently fragmented using High Energy Collisional Dissociation fragmentation, and reporter ions were detected in the Orbitrap. The resolution was set to 60,000 with lower threshold of 110 m/z. AGC of 1 × 10^5^ was used here, with maximum inject time of 100 ms. All data were collected in “centroid” mode.

### Proteomic Data Analysis

Spectral data were analyzed using Proteome Discoverer 2.1. Spectral matching was performed against a concatenated database of *Mus musculus* reference proteome appended to an HSV-1 reference proteome. SEQUEST-HT was used to match spectra against a target and decoy database generated *in silico* (56). Precursor mass and fragment ion mass tolerances were set to 50 ppm and 0.6 Da, respectively. Trypsin was specified as the enzyme, with two missed cleavages permitted. The length of accepted peptides was specified as 6 to 144 amino acids. Oxidation of methionine residues was specified as a variable modification (+15.995 Da), and TMT labeling at N-termini and on lysine residues (+229.163 Da) and carbamidomethylation of cysteine residues (+57.021 Da) were specified as invariable modifications. False discovery rate of 1% was used to filter spectra at the peptide and protein level. Data were processed by filtering peptide spectral matches (PSMs) to retain only high quality and low ambiguity spectra. Any missing quantitation values within a 10-plex were replaced by a 1 to represent a baseline noise value. PSMs with average quantitation values less than 10 or isolation interference greater than 25 were removed from the dataset. PSMs were summed at the protein level and raw values were normalized against the median bridge channel value.

### Quantification and Statistical Analysis

Statistical tests including Mann-Whitney and Wilcoxon signed-rank tests were implemented in GraphPad Prism and *R* where appropriate, as indicated in the figure legends. Details of statistical analysis method and information including *n*, mean, and statistical significance values are indicated in the corresponding sections of the main text, figure legends, or methods. Experiments were replicated three times unless otherwise specified. A p value of < 0.05 was considered to be statistically significant. *p<0.05, **p<0.01, ***p<0.001, ****p<0.0001; ns, not significant.

## Acknowledgments

This work was supported by NIH grants R01EY024710 (DS), R01EY029426 (DS), R01AI139768 (DS), R21AI128171 (DS), R21AI133557 (DS), NIH fellowship F30EY025981 (AA), and departmental core grant P30EY001792 (DS). We are grateful to Ruth Zhelka and Tara Nguyen for assistance with departmental imaging and animal facilities.

## Author Contributions

Conceptualization, A.A. and D.S.; Methodology, A.A., B.A.T., R.K.S., T.Y., D.J., J.A., J.H., L.K., C.D.P., S.R.H., E.J.K., A.C., J.M.W., D.J.G., I.V., J-P.L., D.L.P., P.W.F., D.S.; Software, A.A., B.A.T., A.C., J.M.W.; Validation, A.A., B.A.T., R.K.S., T.Y., D.J., J.A., J.H., L.K., C.D.P., S.R.H., E.J.K., A.C., J.M.W.; Formal Analysis, A.A., B.A.T., A.C., J.M.W.; Investigation, A.A., B.A.T., R.K.S., T.Y., D.J., J.A., J.H., L.K., C.D.P., S.R.H., E.J.K., A.C., J.M.W.; Resources, A.A., B.A.T., A.C., J.M.W., I.V., J-P.L. D.J.G., P.W.F., D.L.P., D.S.; Data Curation, A.A., B.A.T., A.C., J.M.W.; Writing – Original Draft, A.A.; Writing – Review and Editing, A.A., B.A.T., R.K.S., T.Y., D.J., J.A., J.H., L.K., C.D.P., S.R.H., E.J.K., A.C., J.M.W., D.J.G., I.V., J-P.L., D.L.P., P.W.F., D.S.; Visualization, A.A. and B.A.T.; Supervision, A.A., P.W.F., D.L.P., D.J.G, D.S.; Funding Acquisition, A.A. and D.S.

## Supplemental Material Legends

**Figure S1.**
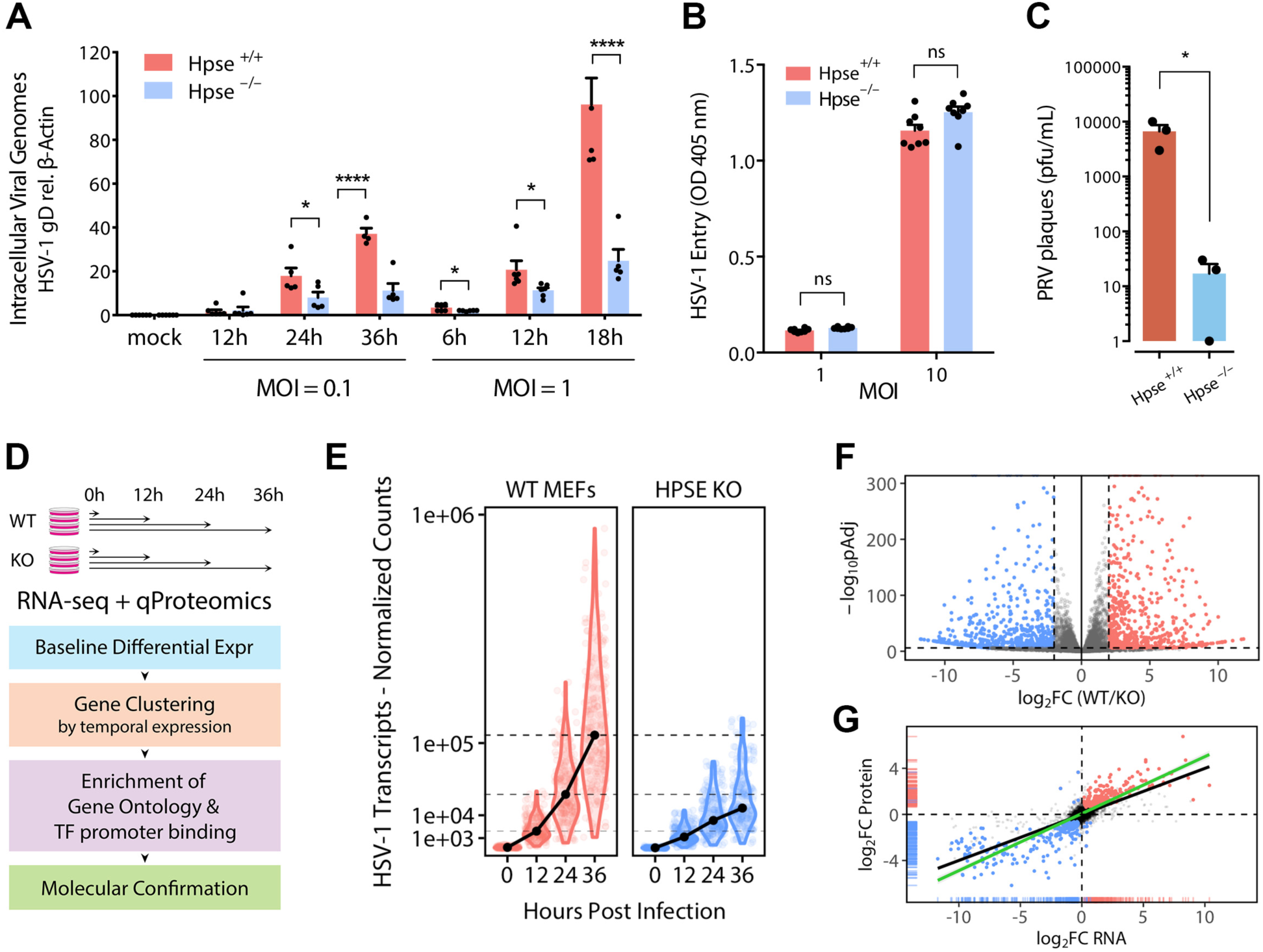
Viral replication is diminished in the absence of HPSE. **(A)** HSV-1 intracellular genome counts relative to β-actin in respective cell type measured by quantitative PCR. **(B)** Viral entry at multiplicity of infection (MOI) of 1 and 10 quantified using β-galactosidase expressing HSV-1 (gL86 strain) and measurement of colorimetric substrate optical density. **(C)** Plaque formation by pseudorabies virus (PRV) in wildtype and Hpse-deficient cells. **(D)** Experimental design and analysis pipeline of unbiased approaches. **(E)** HSV-1 viral gene expression as quantified by transcriptomics analysis. Black dots indicate mean expression for 74 detected viral transcripts. Dashed lines indicate median levels of gene expression in wildtype cells for comparison. **(F)** Volcano plot depicting baseline differential expression in transcriptomic dataset, with red and blue points corresponding to significantly upregulated genes in Hpse ^+/+^ and Hpse ^-/-^ cells, respectively. **(G)** Concordance analysis of baseline RNA-seq and proteomics datasets. Black dots and line indicate non-significantly differentially expressed genes, while colored dots and line represent significantly differentially expressed genes in both datasets. *p<0.05, ****p<0.0001, ns, not significant.

**Figure S2.**
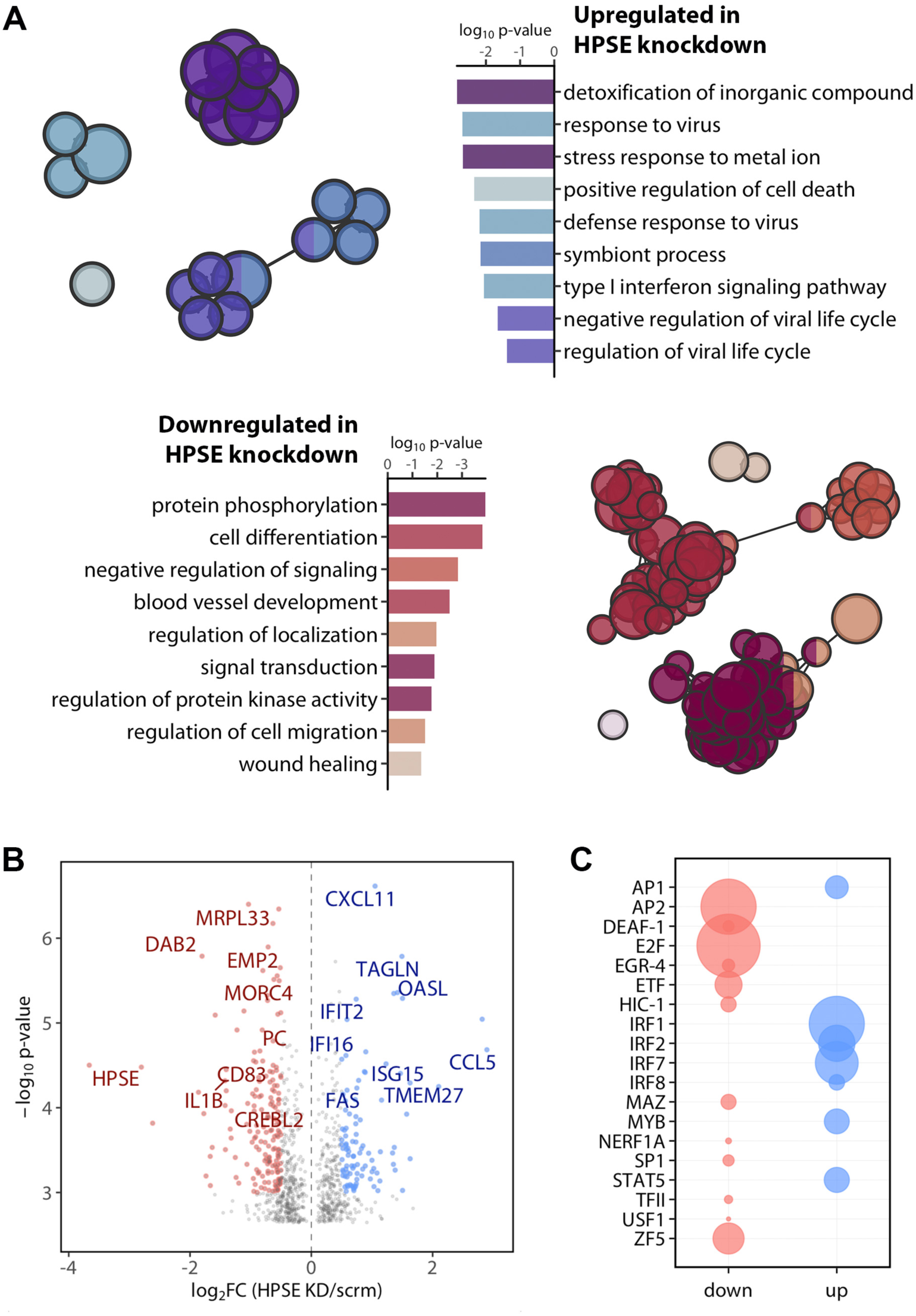
Analysis of publicly available microarray dataset of HPSE knockdown in human cells yields similar results to analysis of HPSE knockout mouse cells. **(A)** Gene ontology (GO) term enrichment analysis of HPSE siRNA knockdown of human gastric carcinoma cells (Gene Expression Omnibus dataset GSE34080), re-analyzed in our lab in R and ClueGO/Cytoscape. Each node represents a significantly enriched, with fold enrichment relative to node size and related terms grouped by color and spatial proximity. Each bar is color coded to match the node it represents. **(B)** Volcano plot indicating gene up- and down-regulation of HPSE-depleted human cells. **(C)** Transcription factor binding site enrichment analysis indicating major factors predicted using the PASTAA algorithm to regulate genes down- or up-regulated in this dataset.

**Figure S3.**
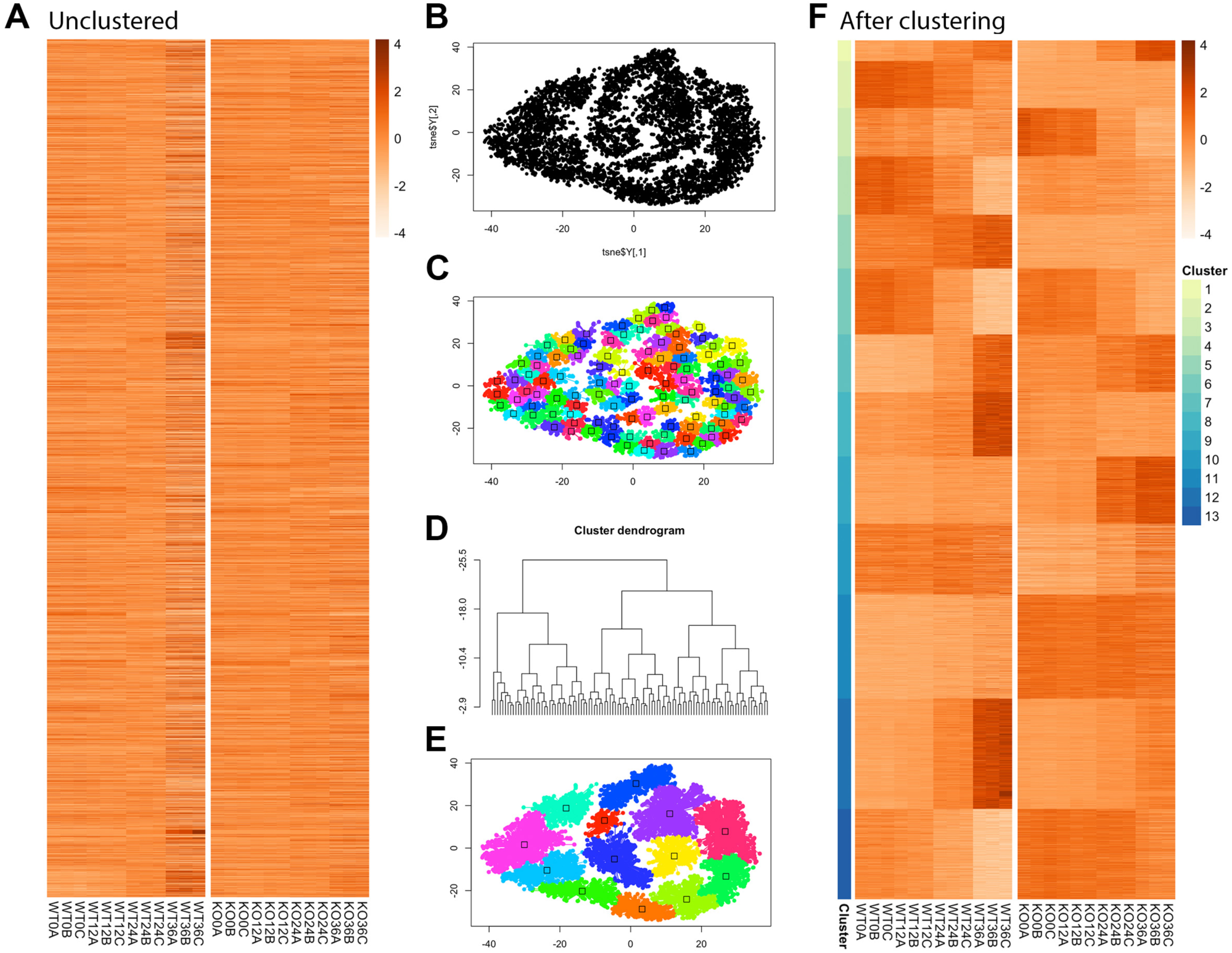
Demonstration of transcriptomics clustering method. **(A)** Heatmap of unclustered significantly differentially expressed genes (*n*=7015) shown in alphabetical order. **(B-E)** Rtsne dimensional reduction and APcluster packages represent all genes in two dimensions and apply clustering based on temporal expression pattern. **(F)** Heatmap of significantly differentially expressed genes represents 13 clearly defined clusters.

**Figure S4.**
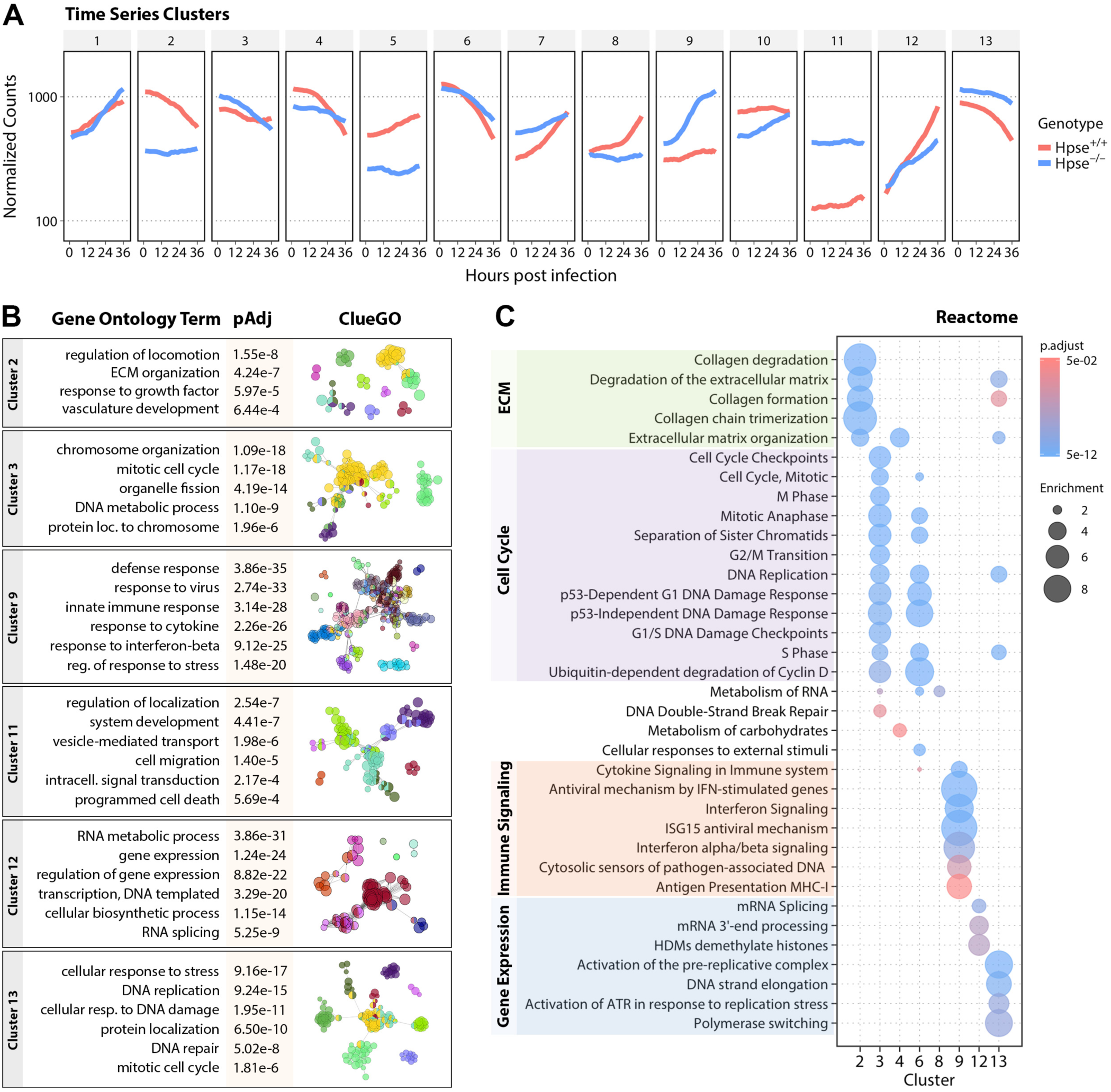
Time series analysis of clusters of significantly differentially expressed genes in HPSE-deficient cells. **(A)** Median gene expression pattern for each of 13 clusters at indicated time post HSV-1 infection. **(B)** Selected gene ontology (GO) results for specified clusters with indicated significance values and diagram of connectedness at right generated with ClueGO/Cytoscape. Each node represents a separate GO term, with node size related to term enrichment and related terms grouped by color and spatial proximity. **(C)** Graphical representation of Reactome analysis indicating major functional enrichments within each gene cluster.

**Figure S5.**
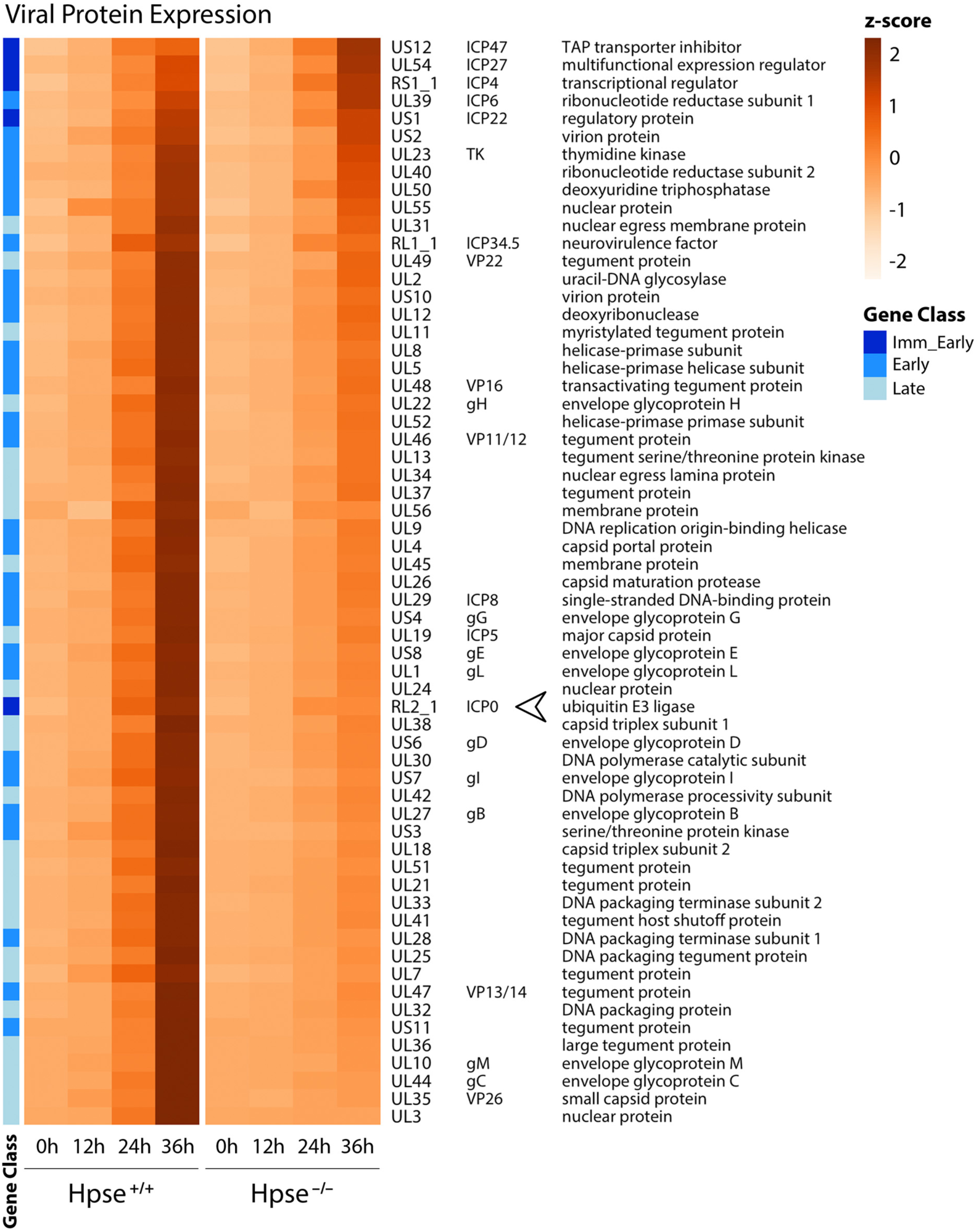
Selective defect in viral protein production in Hpse-deficient cells. Heatmap of all viral proteins detected by quantitative tandem mass tag proteomics analysis. Arrowhead indicates ICP0 as the only immediate early viral protein with markedly diminished levels in Hpse-deficient cells..

